# A Bayesian approach to estimate the probability of resistance to bedaquiline in the presence of a genomic variant

**DOI:** 10.1101/2022.08.30.505812

**Authors:** Degefaye Zelalem Anlay, Emmanuel Rivière, Pham Hien Trang Tu, Steven Abrams, Annelies Van Rie

**Affiliations:** School of Nursing, College of Medicine and Health Science, University of Gondar, Gondar, Ethiopia; Global Health Institute, Family Medicine and Population Health, Faculty of Medicine and Health Sciences, University of Antwerp, Antwerp, Belgium; Data Science Institute, Interuniversity Institute for Biostatistics and statistical Bioinformatics, UHasselt, Diepenbeek, Belgium

**Keywords:** Bayesian, bedaquiline, genomic variant, resistance, probability, Mycobacterium tuberculosis, tuberculosis

## Abstract

**Background:** Bedaquiline is a core drug for treatment of rifampicin-resistant tuberculosis. Few genomic variants have been statistically associated with bedaquiline resistance. Alternative approaches for determining the genotypic-phenotypic association are needed to guide clinical care.

**Methods:** Using published phenotype data for variants in *Rv0678, atpE, pepQ* and *Rv1979c* genes in 756 *Mycobacterium tuberculosis* isolates and survey data of the opinion of 33 experts, we applied Bayesian methods to estimate the posterior probability of bedaquiline resistance and corresponding 95% credible intervals.

**Results:** Experts agreed on the role of *Rv0678*, and *atpE*, were uncertain about the role of *pepQ* and *Rv1979c* variants and overestimated the probability of bedaquiline resistance for most variant types, resulting in lower posterior probabilities compared to prior estimates. The posterior median probability of bedaquiline resistance was low for synonymous mutations in *atpE* (0.1%) and *Rv0678* (3.3%), high for missense mutations in *atpE* (60.8%) and nonsense mutations in *Rv0678* (55.1%), relatively low for missense (31.5%) mutations and frameshift (30.0%) in *Rv0678* and low for missense mutations in *pepQ* (2.6%) and *Rv1979c* (2.9%), but 95% credible intervals were wide.

**Conclusions:** Bayesian probability estimates of bedaquiline resistance given the presence of a specific mutation could be useful for clinical decision-making as it presents interpretable probabilities compared to standard odds ratios. For a newly emerging variant, the probability of resistance for the variant type and gene can still be used to guide clinical decision-making. Future studies should investigate the feasibility of using Bayesian probabilities for bedaquiline resistance in clinical practice.

## Introduction

In 2020, tuberculosis (TB), caused by *Mycobacterium tuberculosis (Mtb)*, was the 13^th^ leading cause of death and the second leading infectious killer after COVID-19 (1). The public health threat posed by TB is worsened by the occurrence of drug-resistant TB. In 2019, close to half a million people developed rifampicin-resistant TB (RR-TB) worldwide (2). In the past decade, important progress has been made, including rapid diagnosis of RR-TB and introduction of short all-oral treatment regimens (3).

Bedaquiline (BDQ) is an essential component of the novel short all-oral RR-TB treatment regimens (4). Its use is rapidly increasing, with 90 countries using BDQ in 2018 (5). Soon after its introduction, cases of acquired and primary BDQ resistance and cross-resistance between BDQ and clofazimine (CFZ), another drug frequently used to treat RR-TB, were reported (6-9). To prevent acquisition and transmission of BDQ resistant TB, the use of BDQ should be accompanied with access to BDQ drug susceptibility tests (DST). Unfortunately, phenotypic DST (pDST) for BDQ, the gold standard, is slow, not yet standardized, and has a low positive predictive value in settings with low prevalence of BDQ resistance (10). A genotypic DST (gDST) assay could provide accurate and timely information if knowledge on genetic variants and their association with resistance to BDQ is comprehensive (11-13).

Several candidate BDQ resistance genes have been identified (14). Mutations in the *atpE* gene can cause a loss of binding affinity of BDQ with its drug target, resulting in high-level BDQ resistance *in vitro* (15), but they are rarely observed in clinical isolates (16). Overexpression of the MmpS5/MmpL5 efflux system caused by mutations in the *Rv0678* gene is the main mechanism of clinical resistance to BDQ (6, 11, 17). Lastly, mutations in the *pepQ* and *Rv1979c* genes have also been implicated in BDQ resistance, but evidence is lacking (9, 18).

A recent systematic review analyzed the association between phenotypic BDQ resistance and genomic variants in *atpE, Rv0678, Rv1979c*, and *pepQ* (13). When applying standard statistical methods, only two out of 313 variants reported in the literature could be statistically associated with resistance (19). Similarly, no mutations satisfied the criteria for BDQ resistance in the 2021 WHO catalogue of mutations in *Mtb* complex (14). Consequently, the body of evidence on BDQ genotype-phenotype association cannot inform the use of gDST for patient care.

Bayesian methods can overcome data sparsity by combining data with expert opinion (20). A Bayesian approach could be of great value in a clinical setting as it is conceptually similar to how clinicians approach the diagnostic process. Furthermore, the credible interval is well suited for clinical practice as it has a more intuitive interpretation (21, 22). We applied a Bayesian method to estimate the probability of BDQ resistance in the presence of genomic variants.

## Materials and Methods

Bayesian methods were used to estimate the probability of BDQ resistance in the presence of a mutation. First, a survey was conducted to obtain expert opinion (prior information) on genotype-phenotype associations for BDQ. Next, an update of a recent systematic literature review was performed to obtain the most comprehensive phenotypic-genotypic data (13). Finally, Bayesian analyses were used to obtain the posterior distribution of the probability of BDQ resistance, the posterior median and its 95% credible interval (CrI) (23).

### Survey

Experts were selected based on their active involvement in the field of genotype-phenotype associations in *Mtb* as determined by scientific publications or participation in the development of the WHO catalogue of mutations in *Mtb* (14). Experts were asked to share contact details of relevant colleagues. The online “Bedaquiline resistance survey” tool (see Survey, Supplementary questionnaire, which includes the consent form and survey presented to experts) focused on the expert rules introduced by *Miotto et al*. (19) and *Köser et al*. (24). For the *atpE, Rv0678, Rv1979c*, and *pepQ* genes, experts were asked if they believe that a mutation can cause BDQ resistance. If the expert answered yes, their opinion on the probability of resistance in the presence of different types of mutations (in-frame indel, frameshift indel, synonymous, nonsense, missense, and homoplastic mutation) was obtained using a six-point Likert scale, except for loss of function mutations (nonsense and frameshift mutation) in the *atpE* gene as these have not been observed in *Mtb* isolates. The survey was piloted by five experts.

### Systematic literature review

To obtain the most comprehensive data on BDQ genotype-phenotype, the individual isolate systematic review (publications between 2008 and October 2020) was updated using the same search engines (Europe PubMed Central and Scopus), search terms and methodology for the period of November 2020 to December 30, 2021 (13). to identify studies that reported genotypic and phenotypic bedaquiline resistance for individual clinical or non-clinical *Mtb* isolates. New records of individual isolates were added to the review database (see Supplementary data, which includes the screened literature, all extracted data, the meta-analysis, expert survey responses, and Bayesian analysis).

### Statistical analysis

All analyses were done using R statistical software (version 4.0.5) (see Supplementary R code).

### Construction of prior distributions

Experts’ responses were eligible for inclusion in the analysis if the expert responded “yes” to “Do you believe that a mutation in the *atpE* gene can confer resistance to BDQ*?”*, indicated he/she was “sure” or “relatively sure” about their responses, and if there was no evidence of poor response quality. The correlation between responses for the same mutation types in different genes was assessed using the Spearman correlation coefficient (25).

Experts’ responses on probability of BDQ resistance in the presence of a variant were translated into prior distributions. For the genes where all experts agreed on its potential role in BDQ resistance, a beta distribution (with shape parameters denoted as α and β) was used to parametrically model the prior distribution for the probability of BDQ resistance for each type of mutation (synonymous mutation, nonsense mutation, frameshift indel, missense mutation, inframe indel, homoplastic mutation). In order to obtain maximum likelihood (ML) estimates for the α- and β-parameters of the prior distribution for each mutation type, the likelihood function was maximized using the *optim* function (in the *stats* R package) with default Nelder-Mead line search method (26). To deal with the interval-censored nature of the Likert scale data (i.e., *x*_*i*_ = [*ll*_*i*_, *ul*_*i*_), for *i= 1*, …, *n*), the (log)likelihood function, denoted hereunder as *LL(x* | α, β), for the estimation of the beta density was adjusted as follows (27):

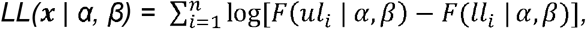

where *x=* (*x*_*1*_, *…, x*_*n*_) represents the vector of reported probability intervals of BDQ resistance, given the presence of a specific type of mutation, for all experts and *F*(*s* | *α, β*) is the cumulative distribution function of a beta distributed random variable with parameters α and β evaluated in s, and *ul_i_* and *ll_i_* the upper and lower limits, respectively, of the probability interval associated with the Likert scale category chosen by expert *i*.

For genes where experts expressed uncertainty about the role in BDQ resistance, a mixture distribution was used as prior. When all three answers (“yes”, “no”, “I do not know”) were observed for the question “can mutations in gene **X** confer resistance to BDQ”, a three-component mixture prior distribution was defined using empirical probability weights *w*_*j*_ :

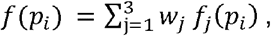

where *f*_*j*_ *(p*_*i*_) represents the probability density for mixture distribution *j* = 1, 2, 3 and nonnegative weights *w*_*j*_ such that 0 < *w*_*j*_ < 1 for all *j* and 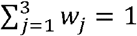 (28). More specifically, f_l_ is the probability density function (pdf) of a beta distribution with shape parameters α_l_ and β_*l*_, *f*_*2*_ is the discrete probability distribution of a degenerate distribution at zero (i.e., *f*_*2*_ (*p*_*i*_) = *Pr*(*P*_*i*_ *= p*_*i*_) = *l*(*p*_*i*_ *= 0*) for *0 < p*_*i*_ *<* 1, where I(.) is the indicator-function) and f_3_ is the pdf of a uniform distribution on the unit interval, which is the same as a beta distribution with *α*_*3*_ = β_3_ =1 (29). The mixture proportions of each component were set equal to the observed probability of responding “yes”, “no” or “I do not know”, respectively.

When only two answers were observed, a two-component mixture prior distribution was used:

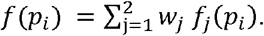

To capture the experts’ opinion on the assumption that the effect of mutations on the phenotype that occurs in laboratory experiments can be extrapolated to the phenotype of clinical isolates containing the same variant, a single unconditional prior distribution was constructed combining the information from experts answering “yes” and the uncertainty from experts answering “I don’t know” or “no.” (30): More specifically, 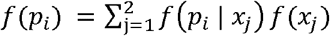 where *f*(*p*_*i*_ | *x*_*j*_) represents the conditional distribution of (*P*_*i*_ | *x* = *x*_*j*_), *j* = 1, 2, for experts answering “yes” (*j* = 1) and “I don’t know” or “no” (*j* = 2), respectively. Finally, *f* represents the discrete probability distribution for the aforementioned response categories, i.e., *f*(*x*_*j*_)= *Pr*(*x* = *x*_*j*_).

### Data likelihood

The genotype-phenotype individual isolate analysis of the updated systematic literature was used for the construction of the data likelihood. Observing phenotypic BDQ resistance (*z*_*k*_ = 1) or susceptibility (*z*_*k*_ = 0) in the presence of a specific mutation (for *k* = 1, …, m) can be viewed as a sequence of independent Bernoulli trials with the same probability of resistance, thereby implying that the number of resistant isolates, denoted by 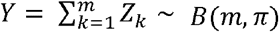, follows a binomial distribution with m the number of times a specific genetic variant in the target gene of interest was reported in the literature, and n the probability of phenotypic BDQ resistance of an isolate containing such variant.

### Posterior distribution and Bayesian inference

In order to estimate the probability of BDQ resistance and its 95% CrI, inferences were made based on the posterior distribution. Dependent Markov Chain Monte Carlo (MCMC) sampling of the posterior is performed using a Metropolis-Hastings (MH) algorithm. Convergence of the MCMC chains was checked by inspection of the trace plots and formal Gelman-Rubin convergence statistics, denoted by R. Three chains with starting values 0.25, 0.5 and 0.75 for the resistance probability were run and convergence was declared when R was less than 1 (31-33). An autocorrelation plot was used to evaluate the dependence of the parameter to the Markov process after the burn-in period.(34) Thinning of the chain was performed every 5^th^, 10^th,^ or 15^th^ iteration, depending on the results of the autocorrelation plot, and the iteration number was increased after thinning. The sufficiency of the iteration number to estimate the 2.5% and 97.5 % quantiles was checked by the Raftery diagnostic (35, 36).

## RESULTS

### Expert survey results and estimation of the prior

Of the 120 invited researchers, 46 (38.3%) completed the survey. Thirteen (28.3%) respondents were excluded from the analysis because they did not fulfill the predefined inclusion criteria. Of the 33 experts whose results could be included, most (n=25, 75.8%) were laboratory researchers and almost all (n=27, 81.8%) resided in a high-income country (Table 1). Regarding general expert rules on genotype-phenotype associations, most experts believed that resistance-conferring mutations occurring in isolation also confer resistance when occurring in combination with other variants (n=26, 78.8%), that any increase above the critical concentration is clinically relevant (n=20, 60.6%), and that the phenotypic effect of a variant in a laboratory strain can be extrapolated to the effect in a clinical isolate (n=22, 66.7%) (Table 2).

**TABLE 1:**
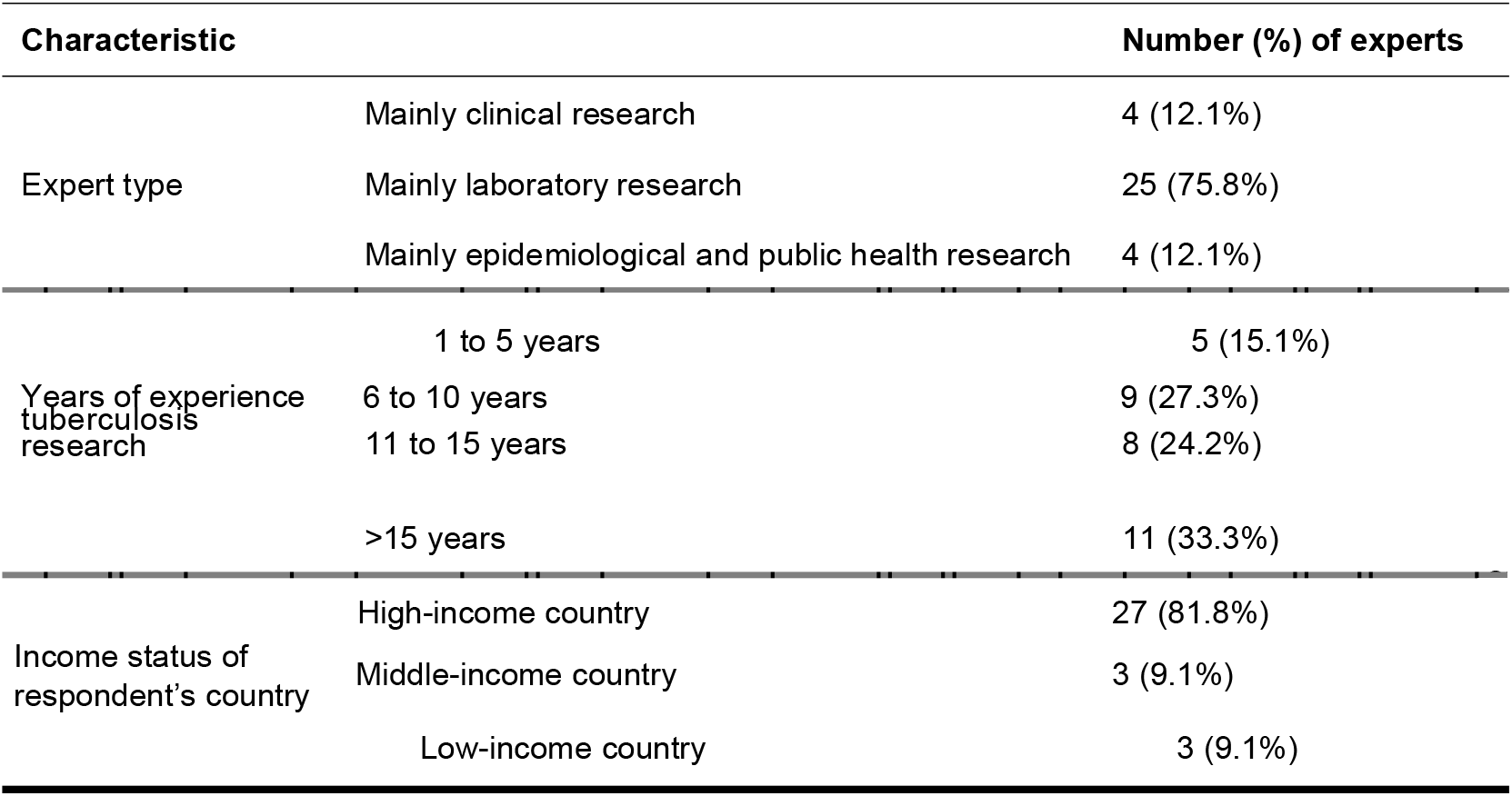
Characteristics of experts contributing data to the expert opinion analysis (n=33)

**TABLE 2:** Expert (n=33) opinion on general rules regarding genotype-phenotype associations for BDQ resistance in *Mtb*

| General expert rule [*Miotto et al.* (19) and *Köser et al.* (24)] | Response | Frequency (%) |
| --- | --- | --- |
| **Rule 1.** A mutation that confers phenotypic resistance to BDQ when it occurs on its own also causes resistance when it occurs in combination with other mutations in the same gene or mutations elsewhere in the genome. | Always | 3 (9.1%) |
|  | Most of the time | 23 (69.7%) |
|  | Sometimes | 6 (18.2%) |
|  | Never | 0 |
|  | I do not know | 1 (3.0%) |
| **Rule 2.** The critical concentration (cc) for BDQ (as defined by WHO or EUCAST) corresponds to a clinical breakpoint, and any minimal inhibitory concentration increase above the cc is clinically significant (i.e., both low level and high-level resistance are clinically relevant for BDQ). | Yes | 20 (60.6%) |
|  | No | 3 (9.1%) |
|  | I do not know | 10 (30.3%) |
| **Rule 3.** The effect of a specific mutation on the phenotype observed in laboratory experiments can be extrapolated to the effect that same mutation would have in clinical isolates. Results of laboratory experiments can, therefore, be used as a source of evidence. | Yes | 22 (66.7%) |
|  | No | 9 (27.3%) |
|  | I do not know | 2 (6.0%) |
| BDQ = bedaquiline; Mtb = Mycobacterium tuberculosis |  |  |

Regarding the role of specific genes in BDQ resistance, all experts believed that a variant in *atpE* or *Rv0678* can cause resistance to BDQ. For the *pepQ* and *Rv1979c* genes, 51.5% (17/33) and 78.1% (25/32) were uncertain whether a variant can cause resistance to BDQ, respectively. The probability that a synonymous mutation can confer resistance was assumed low (never or rarely) for all genes by most experts (*atpE* (84.4%), *Rv0678* (68.8%), *pepQ* (90.0%), *Rv1979c* (100.0%)) (Table 3). Most experts believed that a nonsense mutation frequently to always confers resistance (*Rv0678* (84.8%), *pepQ* (60.0%), *Rv1979c* (50.0%)) and that frameshift indels frequently to always confer resistance when occurring in *Rv0678* (93.8%) or *pepQ* (50.0%) but not when occurring in *Rv1979c* (0.0%). Opinions on the probability of resistance in the presence of a missense mutation or inframe indel varied. The opinion of individual experts on the role of types of mutations in conferring BDQ resistance was correlated for the *atpE* and *Rv0678* genes (synonymous mutations (r=0.38, p=0.03), missense mutations (r=1, p<0.001)).

**TABLE 3:** Expert opinion on the probability of BDQ resistance given the presence of a specific class of mutations

| Mutation type | Probability of phenotypic BDQ resistance |  |  |  |  |  |  |
| --- | --- | --- | --- | --- | --- | --- | --- |
|  | Count<sup>a</sup> | Rarely<br>(<5%) | Occasionally<br>(5-24%) | Sometimes<br>(25-49%) | Frequently<br>(50-74%) | Very frequently<br>(75-94%) | Almost always<br>(≥95%) |
| ***atpE*** |  |  |  |  |  |  |  |
| Synonymous | 32 | 27 | 4 | 1 | 0 | 0 | 0 |
| Missense | 30 | 2 | 4 | 5 | 8 | 7 | 4 |
| Inframe indel | 31 | 5 | 4 | 10 | 5 | 5 | 2 |
| Homoplasic | 33 | 4 | 3 | 3 | 4 | 10 | 9 |
| ***Rv0678*** |  |  |  |  |  |  |  |
| Synonymous | 32 | 22 | 2 | 2 | 3 | 0 | 3 |
| Nonsense | 33 | 0 | 1 | 4 | 5 | 10 | 13 |
| Frameshift | 32 | 1 | 1 | 0 | 8 | 13 | 9 |
| Missense | 30 | 2 | 4 | 5 | 8 | 7 | 4 |
| Inframe indel | 30 | 3 | 2 | 11 | 9 | 5 | 0 |
| Homoplasic | 33 | 1 | 3 | 8 | 5 | 8 | 8 |
| ***pepQ*** |  |  |  |  |  |  |  |
| Synonymous | 10 | 9 | 0 | 0 | 1 | 0 | 0 |
| Nonsense | 10 | 1 | 2 | 1 | 2 | 2 | 2 |
| Frameshift | 10 | 2 | 1 | 2 | 2 | 1 | 2 |
| Missense | 10 | 0 | 4 | 4 | 1 | 1 | 0 |
| Inframe indel | 9 | 3 | 1 | 2 | 2 | 1 | 0 |
| Homoplasic | 10 | 1 | 1 | 3 | 1 | 2 | 2 |
| ***Rv1979c*** |  |  |  |  |  |  |  |
| Synonymous | 2 | 2 | 0 | 0 | 0 | 0 | 0 |
| Nonsense | 2 | 0 | 1 | 0 | 1 | 0 | 0 |
| Frameshift | 2 | 0 | 1 | 1 | 0 | 0 | 0 |
| Missense | 2 | 0 | 2 | 0 | 0 | 0 | 0 |
| Inframe indel | 1 | 1 | 0 | 0 | 0 | 0 | 0 |
| Homoplasic | 2 | 0 | 0 | 1 | 0 | 0 | 1 |

<sup>a</sup> Count For *atpE* and *Rv0678*, all 33 experts believed these genes can play a role in BDQ resistance, but some experts did not complete all questions; for *pepQ* 12 experts believed this gene can play a role but 2 to 3 experts did complete all questions; for *Rv1979c*, only 2 experts believed this gene can play a role in BDQ resistance. BDQ = bedaquiline

The prior median probability of BDQ resistance was lowest in the presence of a synonymous mutation (0.1% for *atpE*, 0.2% for *Rv0678*, 26.0% for *pepQ*), higher for inframe indels (32.8% for *pepQ*, 39.6% for *atpE*, 45.9% for *Rv0678*) and missense mutations (38.0% for *pepQ*, 60.9% for *atpE*, 66.9% for *Rv0678*) and highest for frameshift (46.1% for *pepQ*, 85.2% for *Rv0678*) and nonsense mutations (47.9% for *pepQ*, 90.0% for *Rv0678*). For the *Rv1979c* gene, the prior median probability of resistance was 42.0% for all types of mutations (Table 4; supplementary Tables S1 and S2, showing more shape parameters).

**TABLE 4:**
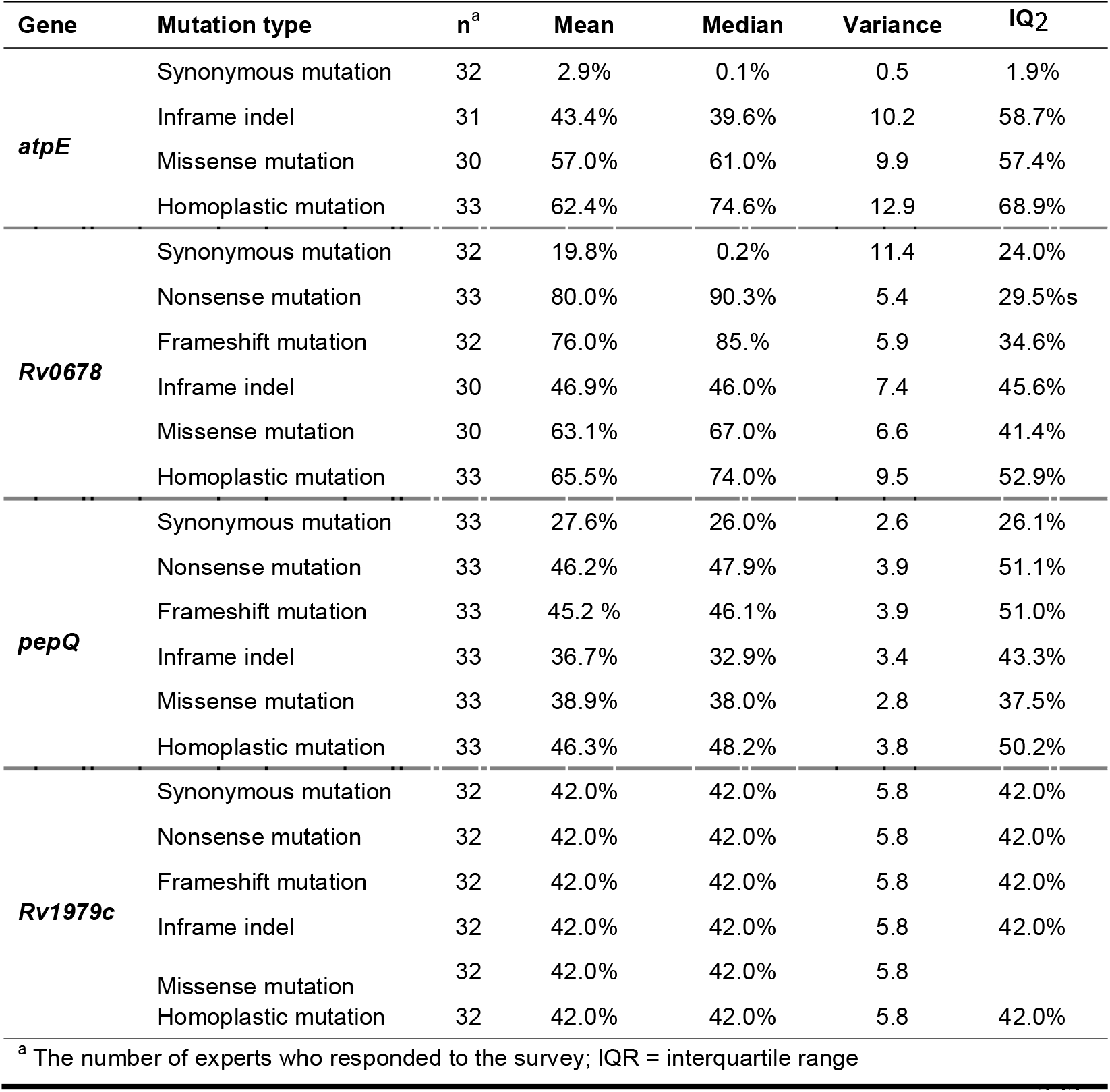
Prior probability distribution for different types of variants by gene of interest

### Updated systematic literature review results

The updated search identified 503 additional studies of which five studies (37-41) were eligible for inclusion in the analysis, bringing the total number of included studies to 46 (see supplementary Figure S1, showing the literature update flowchart). In these 46 studies, 313 unique variants (223 from the published systematic review and 90 additional unique variants) were reported in 756 isolates. Most variants (92.3%, n=289) occurred in clinical isolates of which 98 were reported in isolates phenotypically resistant to BDQ. Of the 189 (25.0%) BDQ resistant isolates, 142 (75.1%) were clinical isolates. Among these 142 phenotypically resistant clinical isolates, the presence of one or more *Rv0678* variant was reported in 106 (74.6%) isolates. The frequency of unique variants ranged from 1 to 40, with *Rv0678* 192_193insG being the most frequent variant (Table 5, 6, 7).

**TABLE 5:** Posterior median probability of BDQ resistance given a specific single nucleotide polymorphism in the *Rv0678* gene

| Rv0678 SNPs | pDST |  |  |  |  |  | OR estimate<br>(95% CI) | Posterior probability<br>estimate (%) (95% CrI) |
| --- | --- | --- | --- | --- | --- | --- | --- | --- |
|  | All |  | Clinical |  | Non-clinical |  |  |  |
|  | R | S | R | S | R | S |  |  |
| C400T (nonsense) | 3 | 0 | 1 | 0 | 2 | 0 | ∞ (1.6-∞) | 96.9 (60.6-99.9) |
| G61T, T276A (nonsense) | 1 | 0 | 0 | 0 | 1 | 0 | ∞ (0.1-∞) | 94.2 (39.2-99.9) |
| C189A | 3 | 0 | 0 | 0 | 3 | 0 | ∞ (1.6-∞) | 82.9 (46.0-99.6) |
| A152C, C158G, G287A | 2 | 0 | 2 | 0 | 0 | 0 | ∞ (0.7-∞) | 84.2 (37.2-99.4) |
| G361A | 3 | 0 | 3 | 0 | 0 | 0 | ∞ (1.6-∞) | 82.9 (46.0-99.6) |
| C403G, G281A | 2 | 0 | 0 | 0 | 2 | 0 | ∞ (0.7-∞) | 84.2 (37.2-99.4) |
| A97G | 2 | 0 | 1 | 0 | 1 | 0 | ∞ (0.7-∞) | 84.2 (37.2-99.4) |
| C148T | 1 | 0 | 0 | 0 | 1 | 0 | ∞ (0.1-∞) | 78.1 (24.6-99.3) |
| C280T | 1 | 0 | 1 | 0 | 0 | 0 | ∞ (0.1-∞) | 78.1 (24.6-99.3) |
| A271C, A413G, A65T, G120C, G197A, T407C | 1 | 0 | 0 | 0 | 1 | 0 | ∞ (0.1-∞) | 78.1 (24.6-99.3) |
| A263G, C155T, C257T, G215A, G326C, T124C,<br>T332A A199C, C247T, C286G, G203A, T118A | 1 | 0 | 1 | 0 | 0 | 0 | ∞ (0.1-∞) | 78.1 (24.6-99.3) |
| T200G | 2 | 0 | 2 | 0 | 0 | 0 | ∞ (0.7-∞) | 84.2 (37.2-99.4) |
| C286T | 2 | 0 | 2 | 0 | 0 | 0 | ∞ (0.1-∞) | 84.2 (37.2-99.4) |
| G5T | 2 | 1 | 2 | 1 | 0 | 0 | 7.6 (0.4-450.5) | 67.1 (25.9-94.5) |
| T157C | 5 | 1 | 5 | 1 | 0 | 0 | 19.3 (2.1-910.4) | 79.7 (45.7-96.9) |
| T131C | 1 | 1 | 0 | 0 | 1 | 1 | 0.0 (0.0-206.3) | 58.7 (16.1-92.8) |
| C403T, A202G | 1 | 1 | 0 | 1 | 1 | 0 | 3.8 (0.1-298.8) | 58.7 (16.1-92.8) |
| C107T, C158T, C176T, C185T, C296T, G193A, T59G,<br>T323C, C143T, T2C | 1 | 1 | 1 | 1 | 0 | 0 | 3.8 (0.1-298.8) | 58.7 (16.1-92.8) |
| C343T(nonsense) | 0 | 1 | 0 | 1 | 0 | 0 | 0.0 (0.0-148.0) | 53.9 (7.3-95.5) |
| C214T | 1 | 2 | 0 | 2 | 1 | 0 | 1.9 (0.0-36.9) | 46.2 (11.5-83.9) |
| A152G | 2 | 2 | 2 | 2 | 0 | 0 | 3.8 (0.3-52.8) | 55.6 (19.2-87.9) |
| A14G, A331G, C-11A, C-53A, C279A, C406G, G109A,<br>G149C, G194C, G196T, G245A, G259A, G352A,<br>G358A, G393C, G421C, G61A, T-20A, T240G, T254G,<br>T274G, T437G, T469G, T93G, A223G, A322G, A44G,<br>A442C, C302A,C319T, C40T, C41T, C474A, C86T,<br>G149A, G184A, G266T, G326A, G467A,T154C | 0 | 1 | 0 | 1 | 0 | 0 | 0.0 (0.0-148.0) | 44.2 (5.6-89.7) |
| T239G | 2 | 2 | 2 | 2 | 0 | 0 | 3.8 (0.3-52.8) | 55.6 (19.2-87.9) |
| G287T | 1 | 1 | 1 | 1 | 0 | 0 | 0.0 (0.0-148.0) | 58.7 (16.1-92.8) |
| A208G | 2 | 1 | 2 | 1 | 0 | 0 | 7.6 (0.4-450.5) | 67.1 (25.9-94.5) |
| T59C | 0 | 2 | 0 | 2 | 0 | 0 | 0.0 (0.0-20.3) | 32.7 (4.0-78.2) |
| T416C | 1 | 1 | 1 | 1 | 0 | 0 | 3.8 (0.1-298.8) | 58.7 (16.1-92.8) |
| C265T (silent) | 1 | 1 | 1 | 1 | 0 | 0 | 3.8 (0.1-298.8) | 43.1 (2.4-94.4) |
| G337A | 1 | 3 | 1 | 3 | 0 | 0 | 1.3 (0.0-15.8) | 38.6 (9.3-76.5) |
| T78G, C112T(nonsense) | 0 | 2 | 0 | 2 | 0 | 0 | 0.0 (0.0-20.3) | 37.6 (4.4-83.5) |
| A-4T, A172C, C247G, G253T, G326T, G362A, G7A,<br>T236C, T278C, G417A, G61A,G250A, T119C | 0 | 2 | 0 | 2 | 0 | 0 | 0.0 (0.0-20.3) | 32.7 (4.0-78.2) |
| G120C | 0 | 2 | 0 | 1 | 0 | 1 | 0.0 (0.0-20.3) | 32.7 (4.0-78.2) |
| G106A, T365C | 0 | 2 | 0 | 2 | 0 | 0 | 0.0 (0.0-20.3) | 32.7 (4.0-78.2) |
| A187G, T425G, A67G, C220G, C251T | 0 | 2 | 0 | 2 | 0 | 0 | 0.0 (0.0-20.3) | 32.7 (4.0-78.2) |
| T248C | 1 | 2 | 1 | 2 | 0 | 0 | 1.9 (0.0-36.8) | 46.2 (11.5-83.9) |
| T136C | 1 | 2 | 1 | 2 | 0 | 0 | 1.9 (0.0-36.8) | 46.2 (11.5-83.9) |
| T350G | 2 | 5 | 2 | 5 | 0 | 0 | 1.5 (0.1-9.4) | 36.9 (11.6-67.8) |
| C6A, G485A, G307A | 0 | 3 | 0 | 3 | 0 | 0 | 0.0 (0.0-9.2) | 26.1 (2.9-68.5) |
| G259C, A292G | 0 | 4 | 0 | 4 | 0 | 0 | 0.0 (0.0-5.8) | 21.7 (2.4-61.3) |
| T341C | 1 | 4 | 1 | 4 | 0 | 0 | 1.0 (0.0-9.7) | 32.8 (7.5-68.4) |
| C268T | 2 | 8 | 2 | 8 | 0 | 0 | 1.0 (0.1-4.8) | 27.6 (8.2-54.8) |
| A11C, A165C | 0 | 5 | 0 | 5 | 0 | 0 | 0.0 (0.0-4.2) | 18.2 (2.0-54.4) |
| A67C | 0 | 6 | 0 | 6 | 0 | 0 | 0.0 (0.0-3.2) | 16.3 (1.8-50.0) |
| T254C, T118G | 0 | 7 | 0 | 7 | 0 | 0 | 0.0 (0.0-2.6) | 14.4 (1.6-44.2) |
| T119C | 0 | 8 | 0 | 8 | 0 | 0 | 0.0 (0.0-20.3) | 13.2 (1.4-40.8) |
| T437C | 1 | 12 | 1 | 12 | 0 | 0 | 0.0 (0.0-2.1) | 15.1 (3.2-37.7) |
| C225T, A318G, C127T, C384A C460T, C81T, G168A,<br>G363A, G390C, G408A,G441A,T297G (Silent) | 0 | 1 | 0 | 1 | 0 | 0 | 0.0 (0.0-148.0) | 0.5 (0.0-68.0) |
| C447T, G195A(silent) | 0 | 2 | 0 | 2 | 0 | 0 | 0.0 (0.0-20.3) | 0.5 (0.0-46.0) |
| G108T, T798G (silent) | 0 | 3 | 0 | 3 | 0 | 0 | 0.0 (0.0-9.2) | 0.4 (0.0-34.0) |
| C305T, T299A, A172C, C201G, T56C, G122T,<br>C143A, A380C, A38C | 1 | 0 | 1 | 0 | 0 | 0 | ∞ (0.1-∞) | 78.1 (24.6-99.3) |
| G137A | 0 | 1 | 0 | 1 | 0 | 0 | 0.0 (0.0-148.0) | 44.2 (5.6-89.7) |

BDQ = bedaquiline; SNP = Single Nucleotide Polymorphism; pDST = phenotypic drug susceptibility test, R = resistant, S = susceptible, OR = odds ratio, CI = confidence interval, CrI = credible interval

**TABLE 6:** Posterior median probability of BDQ resistance of *Rv0678* indels

| Rv0678 indels | pDST |  |  |  |  |  | OR estimate<br>(95% CI) | Posterior probability<br>estimate (%) (95% CrI) |
| --- | --- | --- | --- | --- | --- | --- | --- | --- |
|  | All |  | Clinical |  | Non-clinical |  |  |  |
|  | R | S | R | S | R | S |  |  |
| 198_199insG | 3 | 0 | 3 | 0 | 0 | 0 | ∞ (1.6-∞) | 95.7 (57.4-100.0) |
| 139_141insTG, 145-147indel, 16_17delGG,<br>172_173insIS6110, 18_19delGG, 19delG,<br>212delC, 330delA, 107_108insG, 15delC,<br>325_326insG, 464_466insGC | 1 | 0 | 1 | 0 | 0 | 0 | ∞ (0.1-∞) | 91.1 (36.1-99.9) |
| 259_260insG, 272_273insIS6110,<br>334_335insIS6110, 349_350insIS6110,<br>38_39insA, 65_66insIS6110, 94_95insIS6110 | 1 | 0 | 0 | 0 | 1 | 0 | ∞ (0.1-∞) | 91.1 (36.1-99.9) |
| 138_139insG | 4 | 3 | 4 | 3 | 0 | 0 | 5.2 (0.9-35.7) | 62.3 (29.6-88.4) |
| 107delG, 138_140insGG, 138_140insGA,<br>140_141insG, 142_143delCT, 142_143insC,<br>142delC, 15DelG, 176_177delCG, 176_178insGC,<br>192_194insGG, 193_194insG, 214delC,<br>262_263insA, 274_283delTATTTCCGGT,<br>288delG, 318_320insCG, 335delC, 43_44insA,<br>437_438insT, 457delG, 46indel?, 211delG,<br>28_29insT, 291delC, 29delT, 329delC, 382delG,<br>461delT, 492_493insGA | 0 | 1 | 0 | 1 | 0 | 0 | 0.0 (0.0-148.0) | 51.7 (6.6-94.0) |
| 193delG | 2 | 6 | 0 | 6 | 2 | 0 | 1.3 (0.1-7.3) | 34.2 (10.7-65.3) |
| 140_141insC, 291_292insA, 29delG,<br>359_360insG, 464_465insC, 136_137insG,<br>198delG | 0 | 2 | 0 | 2 | 0 | 0 | 0.0 (0.0-20.3) | 36.2 (4.1-82.6) |
| 400delCGACGGCTGCGG (inframe indel) | 0 | 1 | 0 | 1 | 0 | 0 | 0.0 (0.0-148.0) | 29.7 (1.6-82.2) |
| 133_134insTG, 184_185insC,<br>435delT, 465_466insC | 0 | 3 | 0 | 3 | 0 | 0 | 0.0 (0.0-9.2) | 30.8 (3.1-72.3) |
| 137_138insG | 1 | 3 | 1 | 3 | 0 | 0 | 1.3 (0.0-15.9) | 41.4 (10.1-79.6) |
| 139_140insG | 4 | 3 | 4 | 3 | 0 | 0 | 5.2 (0.9-35.4) | 62.3 (29.6-88.4) |
| 192delG, 288delC | 1 | 6 | 1 | 6 | 0 | 0 | 0.0 (0.0-4.2) | 26.6 (5.7-59.4) |
| 274_275insA | 2 | 6 | 2 | 6 | 0 | 0 | 1.3 (0.1-7.2) | 34.2 (10.7-65.3) |
| 434delT | 0 | 4 | 0 | 4 | 0 | 0 | 0.0 (0.0-5.8) | 22.6 (2.6-64.1) |
| 141_142insC | 4 | 19 | 4 | 19 | 0 | 0 | 0.8 (0.2-2.6) | 21.3 (8.4-40.0) |
| 16delG | 1 | 5 | 1 | 5 | 0 | 0 | 0.0 (0.0-4.2) | 30.3 (6.8-65.2) |
| 418_419insG | 0 | 9 | 0 | 9 | 0 | 0 | 0.0 (0.0-1.9) | 11.8 (1.3-39.5) |
| 192_193insG | 3 | 38 | 3 | 38 | 0 | 0 | 0.3 (0.1-1.0) | 10.0 (3.4-20.9) |
| 11_63del, 430_431insGC, 466_467insGC,<br>141_142insTC, 142_143insGATC, 95delT | 1 | 0 | 1 | 0 | 0 | 0 | ∞ (0.1-∞) | 91.1 (36.1-99.9) |
| del, 181_182insG, 150_151insG | 2 | 0 | 2 | 0 | 0 | 0 | ∞ (0.7-∞) | 93.1 (47.9-99.9) |
| 144_145insC | 10 | 0 | 10 | 0 | 0 | 0 | ∞ (8.9-∞) | 98.2 (79.9-100.0) |
| ins_IS6110 | 1 | 0 | 1 | 0 | 0 | 0 | ∞ (0.2-∞) | 91.1 (36.1-99.9) |

BDQ = bedaquiline; pDST = phenotypic drug susceptibility test, R = resistant, S = susceptible, OR = odds ratio, CI = confidence interval, CrI = credible interval

**TABLE 7:**
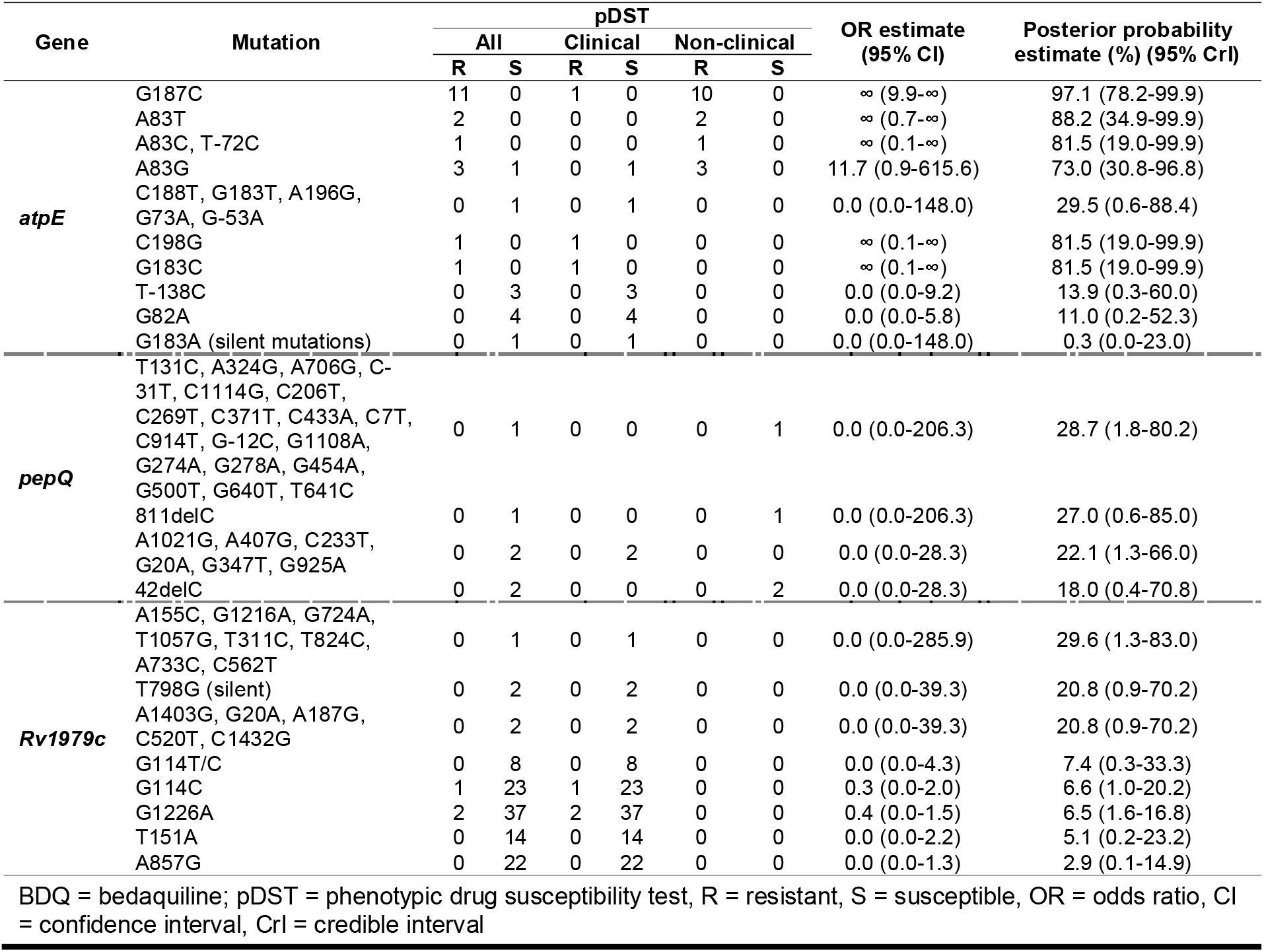
Posterior probability of BDQ resistance given a specific mutation in *atpE, pepQ* and *Rv1979c* genes

### Posterior median probability of BDQ resistance by gene and variant type

For the *atpE* gene, prior (expert opinion) and posterior (prior plus literature review data) probabilities were similar for synonymous mutations (prior median: 0.1%; posterior median: 0.1%) and missense mutations (prior median: 61.0%; posterior median: 60.8%) but 95% CrIs were wide (Table 8). For the *Rv0678* gene, posterior median probabilities were lower than prior median probabilities for frameshift mutations (prior median: 85.2%; posterior median: 30.0%), inframe indels (prior median: 46.0%; posterior median: 29.6%), missense mutations (prior median: 67.0%; posterior median: 31.5%) and nonsense mutations (prior median: 90.3%; posterior median: 55.1%), and similar for synonymous mutations (prior median: 0.2%; posterior median: 3.3%), but all with wide 95% CrIs. For the *pepQ* and *Rv1979c* genes, the posterior median probabilities were lower than the prior median probabilities for frameshift mutations in *pepQ* (prior median: 46.1%; posterior median: 13.7%), missense mutations in *pepQ* (prior median: 26.0%; posterior median: 2.6%), missense mutations in *Rv1979c* (prior median: 42.0%; posterior median: 25.0%) and synonymous mutations in *Rv1979c* (prior median: 42.0%; posterior median: 2.9%), with mostly wide 95% CrIs (Table 8).

**TABLE 8:** Prior and posterior probability of BDQ resistance by mutation type and gene of interest

| Gene | Mutation type | Prior |  | pDST |  | Posterior |  | 95% Crl |
| --- | --- | --- | --- | --- | --- | --- | --- | --- |
|  |  | Mean probability | Median probability | R | S | Mean probability | Median probability |  |
| *atpE* | Synonymous | 2.9% | 0.1% | 0 | 1 | 2.4% | 0.1% | 0.0 - 19.7% |
|  | Missense | 57.0% | 61.0% | 20 | 13 | 60.5% | 60.8% | 44.0 - 76.1% |
| *Rv0678* | Synonymous | 19.8% | 0.2% | 1 | 23 | 4.5% | 3.3% | 0.2 - 15.4% |
|  | Nonsense | 80.0% | 90.3% | 5 | 5 | 54.8% | 55.1% | 27.3 - 80.7% |
|  | Frameshift | 76.0% | 85.2% | 69 | 164 | 30.0% | 30.0% | 24.2 - 36.1% |
|  | Inframe indel | 46.9% | 46.0% | 0 | 1 | 33.0% | 29.6% | 1.7 - 81.8 % |
|  | Missense | 63.1% | 67.0% | 91 | 200 | 31.5% | 31.5% | 26.3 - 36.8% |
| *pepQ* | Frameshift | 45.2% | 46.1% | 0 | 3 | 18.1% | 13.7% | 0.2 - 59.0% |
|  | Missense | 38.8% | 26.0% | 0 | 33 | 3.4% | 2.6% | 0.1 - 11.7% |
| *Rv1979c* | Synonymous | 42.0% | 42.0% | 0 | 2 | 20.8% | 25.0% | 0.9 - 70.2% |
|  | Missense | 42.0% | 42.0% | 3 | 122 | 3.2% | 2.9% | 0.9 - 6.9% |

BDQ = bedaquiline; Crl = Credible interval; pDST = phenotypic drug susceptibility; R = Resistant; S = Susceptible

### Posterior median probability of BDQ resistance for specific mutations

For single nucleotide polymorphisms (SNPs), the probability of resistance ranged from 0.4% for the silent mutations (*Rv0678_*C108T, *Rv0678*_T798G and *atpE*_G183A) to ≥95% for the nonsense mutation *Rv0678_*C400T and missense mutation *atpE*_G187C. However, some missense and nonsense mutations had a substantially lower median probability, while some silent mutations had a much higher probability of resistance (Table 5 and 7). For SNPs, the probability of resistance ranged from 0.3% to 97.1% in *atpE*, from 0.4% to 96.9% in *Rv0678*, from 22.1% to 28.7% in *pepQ*, and from 2.9% to 29.6% in *Rv1979c*. For indels, the probability of resistance ranged from 10.0% to 98.2% in *Rv0678*, and from 18% to 27% in *pepQ* (Table 6 and 7). The 95% CrIs were wide for almost all variants.

## DISCUSSION

Accurate knowledge of the presence of BDQ resistance is important when initiating a BDQ-containing treatment regimen or changing a BDQ-containing regimen when a patient does not respond to their RR-TB treatment. Because pDST requires a lengthy culture process, there is great interest in a genotypic assay. Unfortunately, developing such assays is challenging given the existence of multiple BDQ resistance candidate genes, the occurrence of variants across the genes, and the difficulty to confidently assign a BDQ phenotype to genetic variants using standard statistical methods due to the low prevalence of variants (13, 42, 43).

To overcome these challenges, we combined expert opinions and published genotype-phenotype data to estimate the Bayesian probability of BDQ resistance in the presence of a genomic variant. Our results showed that many experts were uncertain about the role of *pepQ* and *Rv1979c* variants in BDQ resistance and that experts overestimated the probability of resistance for most variant types, resulting in lower posterior (median) probabilities as compared to prior estimates. As expected, the posterior probability of BDQ resistance was low for synonymous mutations in *atpE* and *Rv0678* (<5%) and relatively high for missense mutations in *atpE* (60.8%) and nonsense mutations in *Rv0678* (55.1%). Surprisingly, the posterior probability of BDQ resistance was relatively low for missense mutations and frameshift in *Rv0678* (≤35%) and low for missense mutations in *pepQ* and *Rv1979c* (<3%). For specific SNPs and indels, the probability of resistance varied greatly between variants of the same type, even in the same gene, and 95% CrIs were wide due to sparse data.

For clinicians, the interpretation of a probability may be more intuitive than the interpretation of an odds ratio. For example, for variant *Rv0678*_A202G, the odds ratio of 3.8 (95% CI of 0.05-298.8) is interpreted as: the odds of resistance to BDQ for an *Mtb* isolate with a A202G variant is 3.8 times that of isolates without the A202G variant, independent of the presence of wild type or any other variant, and if we performed the analysis 100 times using different datasets (based on different samples), the estimate of the odds ratio would lie between the lower and upper limit of the constructed CI in 95% of the analyses. In contrast, the interpretation of the probability of resistance to BDQ for an *Mtb* isolate with a A202G variant derived from a posterior distribution with median (or mean) of 58.7% (95% CrI 16.1-92.8%) is more intuitive: the probability of resistance to BDQ for the isolate containing the A202G variant is estimated to be 58.7% and we are 95% certain that the true probability of resistance lies between 16.1% and 92.8%, given the data at hand.

The results of our study should be interpreted in light of some limitations. First, while the most accurate priors would be obtained by limiting the survey to the most knowledgeable experts, we deliberately extended the survey to obtain a broad and globally representative perspective. This could have caused the wide range of opinions and discrepancies between the data and the prior. In the face of discrepancies, determining whether the prior or the data is correct is challenging (23). Second, in the construction of the prior distributions, we did not account for the uncertainty around the estimated shape parameters of the beta distributions.

In order to do so, one would need to specify hyperprior distributions for α and β thereby increasing posterior uncertainty further. Third, the published expert rules were developed for resistant to other drugs used to treat TB and may not fully apply to BDQ. For example, while *Miotto et al* assumed that nonsense and frameshift mutations can be predicted with confidence to cause a loss-of-function phenotype (19, 24), we found that the posterior probability of BDQ resistance in the presence of a nonsense mutation in *Rv0678* was only 55.1% and the probability of resistance for frameshift mutations in *Rv0678* was only 29.9%. Fourth, the phenotype-genotype reported in studies included in the analysis may not always be accurate due to lack of standardization of BDQ pDST. Fifth, to maximally use all reported data on BDQ resistance given a specific mutation, data on clinical and non-clinical isolates was combined and analyzed. Furthermore, the published data may not reflect the actual sampling distributions of the phenotype for a specific variant. Finally, posterior distributions can only be estimated for variants that have been reported in the literature. Given that the introduction of BDQ is recent, it is possible that many variants have not yet been observed or reported in the literature. The strength of the Bayesian approach is that it also provides a resistance probability for different variant types in the candidate resistance genes, which can give guidance to the clinician when confronted with a novel variant.

Probabilistic approaches to predict phenotypic resistance to BDQ from genotypic data could be an alternative approach to the standard statistical methods and help guide treatment decisions as it presents physicians with more intuitively interpretable probabilities (and 95% CrI) as compared to odds ratios (and corresponding 95% CI). For a variant not previously reported, the probability of resistance for a type of variant in a specific gene can still be used to guide clinical decision making. Since a probabilistic approach is new for clinicians, studies should investigate how feasible it is for physicians to use probabilities of BDQ resistance in clinical practice.

## Supporting information

questionnaire

## Ethical approval

The study has been reviewed and approved by the Research Ethics Committee of the University Hospital of the University of Antwerp **(REF number: 21/06/093)**. The survey was anonymous, all participated experts gave written informed consent.

## Data availability

All data extracted for the systematic review, all expert responses and R code are made available as supplementary information.

## Acknowledgements

We thank Claudio Köser for his input into the development of the survey tool and the Tuberculosis Omics ResearCH team members for piloting the survey tool. We also thank the scientists who participated in the survey.

## Author contributions

AVR, ER, DA, and SA conceived the study. ER, AVR, TT, and DA participated in the survey tool development. DA performed the analysis under supervision of SA. The manuscript was written by DA and edited and approved by AVR, ER, TT and SA.

