## supplementary tables for "A Bayesian approach to estimate the probability of resistance to bedaquiline in the presence of a genomic variant"

**Supplementary_table 1:** Prior probablity distribution parametrised by beta shape parameters (α and β) based on experts response in the three of genes of interest

| **Genes** | **Mutation class** | **n*** | $\hat{\boldsymbol{\alpha}}$ **^@^** | **SE(̂**$\hat{\boldsymbol{\alpha}}$**)** | $\hat{\boldsymbol{\beta}}$**^@^** | **SE(**$\hat{\boldsymbol{\beta}}$**)** | **Mean** | **Median** | **Variance** | **IQR** |
| --- | --- | --- | --- | --- | --- | --- | --- | --- | --- | --- |
| ***atpE*** | Synonymous mutation | 32 | 0.135 | 0.005 | 4.541 | 0.377 | 2.9% | 0.1% | 0.5 | 1.9% |
|  | Inframe indel | 31 | 0.607 | 0.023 | 0.793 | 0.032 | 43.4% | 39.6% | 10.2 | 58.7% |
|  | Missense mutation | 30 | 0.840 | 0.034 | 0.630 | 0.025 | 57.0% | 61.0% | 9.9 | 57.4% |
|  | Homoplastic mutation | 33 | 0.508 | 0.022 | 0.305 | 0.011 | 62.4% | 74.6% | 12.9 | 68.9% |
| **Rv0678** | Synonymous mutation | 32 | 0.078 | 0.003 | 0.315 | 0.016 | 19.8% | 0.2% | 11.4 | 24.0% |
|  | Nonsense mutation | 33 | 1.550 | 0.077 | 0.380 | 0.014 | 80.0% | 90.3% | 5.4 | 29.5% |
|  | Frameshift mutation | 32 | 1.550 | 0.076 | 0.480 | 0.018 | 76.0% | 85.2% | 5.9 | 34.6% |
|  | Inframe indel | 30 | 1.112 | 0.042 | 1.257 | 0.053 | 46.9% | 456.0% | 7.4 | 45.6% |
|  | Missense mutation | 30 | 1.596 | 0.067 | 0.932 | 0.037 | 63.1% | 67.0% | 6.6 | 41.4% |
|  | Homoplastic mutation | 33 | 0.904 | 0.039 | 0.476 | 0.018 | 65.5% | 74.0% | 9.5 | 52.9% |
| ***pepQ^#^*** | Synonymous mutation | 10 | 0.027 | 0.001 | 0.584 | 0.066 | 4.4% | 26.0% | 2.6 | 26.1% |
|  | Nonsense mutation | 10 | 0.548 | 0.022 | 0.430 | 0.017 | 56.0% | 47.9% | 12.4 | 51.1% |
|  | Frameshift mutation | 10 | 0.543 | 0.021 | 0.476 | 0.019 | 53.3% | 46.1% | 12.3 | 51.0% |
|  | Inframe indel | 9 | 0.406 | 0.015 | 0.954 | 0.040 | 29.9% | 32.9% | 8.9 | 43.3% |
|  | Missense mutation | 10 | 1.499 | 0.055 | 2.680 | 0.113 | 35.8% | 38.0% | 4.4 | 37.5% |
|  | Homoplastic mutation | 10 | 0.598 | 0.024 | 0.460 | 0.018 | 56.6% | 48.2% | 11.9 | 50.2% |

**n*-**the number of responses varied depending on the belief of the expert whether the gene plays a role in BDQ resistance and the response for a specific type of variant

**#** The mean and the variance is estimated from experts that believe *pepQ* plays a role in BDQ resistance (one of the components in the mixture distribution) and therefore the result presented here is not the final one.

**^@^** Two-thirds of the experts assumed that the effect of mutations on the phenotype that occurs in laboratory experiments can be extrapolated to the phenotype of clinical isolates containing the same variant. A single prior distribution was thus constructed combining the information from experts answering "Yes" (n1=22, p(X1)=0.667) and reflecting the uncertainty coming from experts answering the question about extrapolation as "I don't know" or "No" (n=11, p(X2)=0.333)***.***

**NB:** For Rv1979c the mean prior probablity is not presented in this table because all experts respond “No” or “I don’t know” for the survey question “can mutations in gene Rv1979c confer resistance to BDQ”.

**Supplementary_table 2:** Mixture distribution and mixing proportion for the prior distribution of *pepQ* and Rv1979c

| **Gene** | **Mutation** | **Mixture distribution #** | **Prior median** | **IQR** |
| --- | --- | --- | --- | --- |
| pepQ  *(n=33,*  *yes=12,*  *no=4,*  *I do not know=17)* | S**ynonymous** | f(x)= 0.36*beta(0.027, 0.584) + 0.12*I(x=0) + 0.52*beta (1, 1) | 26.0% | 26.1% |
|  | Nonsense | f(x)= 0.36*beta (0.548, 0.43) + 0.12*I(x=0) + 0.52*beta (1, 1) | 47.9% | 51.1% |
|  | Frameshift | f(x)= 0.36*beta(0.543, 0.475) + 0.12*I(x=0) + 0.52*beta (1, 1) | 46.1% | 51.0% |
|  | Inframe indel | f(x)= 0.36*beta(0.406, 0.954) + 0.12*I(x=0) + 0.52*beta (1, 1) | 32.9% | 43.3% |
|  | Missense | f(x)= 0.36*beta(1.499, 2.688) + 0.12*I(x=0) + 0.52*beta (1, 1) | 38.0% | 37.5% |
|  | Homoplastic | f(x)= 0.36*beta(0.598, 0.460) + 0.12*I(x=0) + 0.52*beta (1, 1) | 48.2% | 50.2% |
| Rv1979c  *(n=32,*  *yes=2,*  *no=5,*  *I don’t know =25)* | S**ynonymous** | f(x)= 0.16*p(x=0) + 0.84*beta (1, 1) | 42.0% | 42.0% |
|  | Nonsense | f(x)= 0.16*p(x=0) + 0.84*beta (1, 1) | 42.0% | 42.0% |
|  | Frameshift | f(x)= 0.16*p(x=0) + 0.84*beta (1, 1) | 42.0% | 42.0% |
|  | Inframe indel | f(x)= 0.16*p(x=0) + 0.84*beta (1, 1) | 42.0% | 42.0% |
|  | Missense | f(x)= 0.16*p(x=0) + 0.84*beta (1, 1) | 42.0% | 42.0% |
|  | Homoplastic | f(x)= 0.16*p(x=0) + 0.84*beta (1, 1) | 42.0% | 42.0% |

**#** The prior distributions for *pepQ* were estimated from three components with a different mixing proportion; The mixture proportions of each component were set equal to the observed probability of responding yes, no or I do not know. “yes” (n1 = 12; $w_{1}=0.36$), “no” (n2 = 4; $w_{2}=0.12$), or “I do not know” (n3 = 17; $w_{3}=0.52$ ) out of the total number included experts (n=33). Whereas for Rv1979c The observed empirical probability of responding “no” (n1 = 5; $w_{1}=0.16$), “I do not know” (n2 = 27; $w_{2}=0.84$), out of the total number included experts (n=32). Two experts who responded "yes" were included in the group who answered "I do not know" because they also disagreed.

**I(.)** is the indicator function being one if the argument is true

IQR: Interquartile range
