## Supplementary figures and images for "A Bayesian approach to estimate the probability of resistance to bedaquiline in the presence of a genomic variant"

### supplementary figures

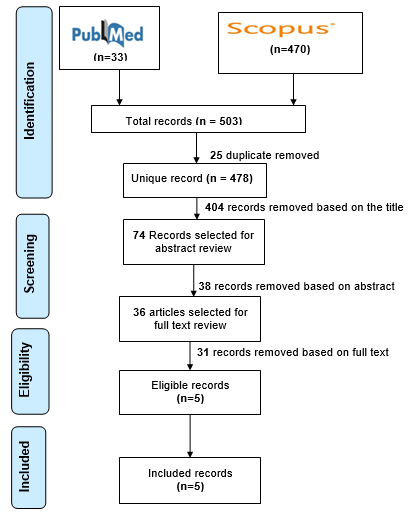

**Figure S1:** Study selection after the published systematic review and individual meta-analysis
