## Supplementary material for "A Bayesian approach to estimate the probability of resistance to bedaquiline in the presence of a genomic variant": questionnaire

**Consent form**

**Study name:** “A Bayesian approach to estimate probabilities of resistance to bedaquiline in the presence of a genomic variant and how this influences the decision-making process of physicians in prescribing bedaquiline”

**Purpose:** We would like to gain insight into your assumptions regarding the probability that a certain class of mutations in different genes of interest will confer resistance to bedaquiline.

**Procedure:** This survey will take approximately 10 minutes of your time

**Voluntary participation:** your participation in this survey is voluntary. You are free to leave the survey at any time.

**Confidentiality:** This survey is anonymous, and all information will remain confidential. Survey responses will be secured on a password protected device and only accessible by the research team. Personal identifiable information will not be accessible to the research team. It will thus not be possible to link your responses to your name.

**Risks:** we do not foresee any risks related to your study participation

**Ethics approval:** this study has been reviewed by the Research Ethics Committee of the University Hospital of the University of Antwerp (REF number: 21/06/093). If you have any questions about your rights as a survey participant, you can contact the ethics committee at

**Researcher contacts:**

-       Annelies Van Rie: 

-       Degafaye Anlay: 

**Consent:** Filling out the survey provides consent for participation in this study.

- **I consent**
- **I do not consent**

***We first want to ask your opinion on three general expert rules.***

**Q1: Expert rule:** A mutation that confers phenotypic resistance to BDQ when it occurs on its own also causes resistance when it occurs in combination with other mutations in the same gene or mutations elsewhere in the genome.

- Always
- Most of the time
- Sometimes
- Never
- I do not know

**Q2: Expert rule**: The critical concentration (cc) for BDQ [as currently defined by WHO or EUCAST] corresponds to a clinical breakpoint and any MIC increase above the cc is clinically significant (i.e. both low level and high level resistance are clinically relevant for BDQ).

- Yes
- No
- I do not know

**Q3: Expert rule**: The effect of a specific mutation on the phenotype observed in laboratory experiments can be extrapolated to the effect that same mutation would have in clinical isolates. Results of laboratory experiments can, therefore, be used as a source of evidence.

- Yes
- No
- I do not know

**Instruction for question (4-7)**

**Information: Studies have suggested that atpE, Rv0678, pepQ, and Rv1979c genes may be involved in BDQ resistance, in vitro and/or in vivo.** 

**In the next section, we ask your opinion on the role of these genes in BDQ resistance. *The answers to some of these questions are currently not known, so there may be no right or wrong answer. We, therefore, ask you to give your expert opinion. If you feel that you don't have enough information to answer for some of the questions you can leave blank for that specific questions and answer others.***

**Q4:** Do you believe that a mutation in ***atpE*** can confer resistance to BDQ?

Yes	No	I do not know	*If* ***yes****, display sub-questions*

**Q4.1: A** **synonymous mutation** [defined as mutation that does not alter the amino acid encoded by the affected codon] **in atpE is neutral** [defined by susceptible phenotype to BDQ]

| Rarely  (<5%) | Occasionally     (5-24%) | Sometimes  (25-49%) | Frequently  (50-74%) | Frequently  (50-74%) | Very frequently     (75-94%) | Almost always (>=95%) |
| --- | --- | --- | --- | --- | --- | --- |

Comment:

**Q4.2: A** **missense mutation** [defined as a mutation that alters the amino acid encoded by the affected codon] **in *atpE* confer resistance to BDQ.**

| Rarely  (<5%) | Occasionally     (5-24%) | Sometimes  (25-49%) | Frequently  (50-74%) | Frequently  (50-74%) | Very frequently     (75-94%) | Almost always (>=95%) |
| --- | --- | --- | --- | --- | --- | --- |

Comment:

**Q4.3: A homoplastic mutation** [defined as a variant that has arisen multiple times independently] **in *atpE* is a likely signal for positive selection and confer resistance to BDQ.**

| Rarely  (<5%) | Occasionally     (5-24%) | Sometimes  (25-49%) | Frequently  (50-74%) | Frequently  (50-74%) | Very frequently     (75-94%) | Almost always (>=95%) |
| --- | --- | --- | --- | --- | --- | --- |

Comment:

**Q4.4: An in-frame indel** [defined as an insertion or deletion of a number - a multiple of three - of nucleotides in a DNA sequence which does not shift the reading frame downstream] **in *atpE* confer resistance to BDQ.**

| Rarely  (<5%) | Occasionally     (5-24%) | Sometimes  (25-49%) | Frequently  (50-74%) | Frequently  (50-74%) | Very frequently     (75-94%) | Almost always (>=95%) |
| --- | --- | --- | --- | --- | --- | --- |

Comment:

**Q5:** Do you believe that a mutation in ***Rv0678*** can confer resistance to BDQ [assuming that the mmpS5/L5 efflux pump is functional]?

Yes	No	*If* ***yes****, display sub-questions*

**Q5.1:** A **synonymous mutation** [defined as mutation that does not alter the amino acid encoded by the affected codon] **in *Rv0678* is neutral** [defined by susceptible phenotype to BDQ]

| Rarely  (<5%) | Occasionally     (5-24%) | Sometimes  (25-49%) | Frequently  (50-74%) | Frequently  (50-74%) | Very frequently     (75-94%) | Almost always (>=95%) |
| --- | --- | --- | --- | --- | --- | --- |

Comment:

**Q5.2: A** **nonsense mutation** [also called a premature stop codon and defined as a substitution of a single base pair that alters the DNA sequence and leads to the production of a shortened protein] **in *Rv0678* is a loss of function mutation and confer resistance to BDQ.**

| Rarely  (<5%) | Occasionally     (5-24%) | Sometimes  (25-49%) | Frequently  (50-74%) | Frequently  (50-74%) | Very frequently     (75-94%) | Almost always (>=95%) |
| --- | --- | --- | --- | --- | --- | --- |

Comment:

**Q5.3:** A **frameshift indel** [defined as an insertion or deletion of a number-not divisible by three- of nucleotides in a DNA sequence that affects the reading frame of the gene and results in a different translation compared to the wild type] **in *Rv0678* is a loss of function mutation and confer resistance to BDQ.**

| Rarely  (<5%) | Occasionally     (5-24%) | Sometimes  (25-49%) | Frequently  (50-74%) | Frequently  (50-74%) | Very frequently     (75-94%) | Almost always (>=95%) |
| --- | --- | --- | --- | --- | --- | --- |

Comment:

**Q5.4: An in-frame indel** [defined as an insertion or deletion of a number - a multiple of three - of nucleotides in a DNA sequence which does not shift the reading frame downstream] **in *Rv0678* confer resistance to BDQ.**

| Rarely  (<5%) | Occasionally     (5-24%) | Sometimes  (25-49%) | Frequently  (50-74%) | Frequently  (50-74%) | Very frequently     (75-94%) | Almost always (>=95%) |
| --- | --- | --- | --- | --- | --- | --- |

Comment:

**Q5.5: missense mutation** other than a nonsense mutation [defined as mutation that alters the amino acid encoded by the affected codon] in *Rv0678* should confer resistance to BDQ.

| Rarely  (<5%) | Occasionally     (5-24%) | Sometimes  (25-49%) | Frequently  (50-74%) | Frequently  (50-74%) | Very frequently     (75-94%) | Almost always (>=95%) |
| --- | --- | --- | --- | --- | --- | --- |

Comment:

**Q5.6: Expert rule:** A **homoplastic mutation** [defined as a variant that has arisen multiple times independently] **in *Rv0678* is a likely signal for positive selection and confer resistance to BDQ.**

| Rarely  (<5%) | Occasionally     (5-24%) | Sometimes  (25-49%) | Frequently  (50-74%) | Frequently  (50-74%) | Very frequently     (75-94%) | Almost always (>=95%) |
| --- | --- | --- | --- | --- | --- | --- |

Comment:

**Question 6:** Do you believe that a mutation in ***pepQ*** can confer resistance to BDQ?

Yes	No	*If* ***yes****, display sub-questions*

**Q6.1:** **A** **synonymous mutation** [defined as mutation that does not alter the amino acid encoded by the affected codon] **in *pepQ*** **is neutral** [defined by susceptible phenotype to BDQ]

| Rarely  (<5%) | Occasionally     (5-24%) | Sometimes  (25-49%) | Frequently  (50-74%) | Frequently  (50-74%) | Very frequently     (75-94%) | Almost always (>=95%) |
| --- | --- | --- | --- | --- | --- | --- |

Comment:

**Q6.2: A** **nonsense mutation** [also called a premature stop codon and defined as a substitution of a single base pair that alters the DNA sequence and leads to the production of a shortened protein] **in *pepQ* is a loss of function mutation and confer resistance to BDQ.**

| Rarely  (<5%) | Occasionally     (5-24%) | Sometimes  (25-49%) | Frequently  (50-74%) | Frequently  (50-74%) | Very frequently     (75-94%) | Almost always (>=95%) |
| --- | --- | --- | --- | --- | --- | --- |

Comment:

**Q6.3: A** **frameshift indel** [defined as an insertion or deletion of a number-not divisible by three- of nucleotides in a DNA sequence that affects the reading frame of the gene and results in a different translation compared to the wild type] **in *pepQ* is a loss of function mutation and confer resistance to BDQ.**

| Rarely  (<5%) | Occasionally     (5-24%) | Sometimes  (25-49%) | Frequently  (50-74%) | Frequently  (50-74%) | Very frequently     (75-94%) | Almost always (>=95%) |
| --- | --- | --- | --- | --- | --- | --- |

Comment:

**Q6.4: An in-frame indel** [defined as an insertion or deletion of a number - a multiple of three - of nucleotides in a DNA sequence which does not shift the reading frame downstream] **in *pepQ* confer resistance to BDQ.**

| Rarely  (<5%) | Occasionally     (5-24%) | Sometimes  (25-49%) | Frequently  (50-74%) | Frequently  (50-74%) | Very frequently     (75-94%) | Almost always (>=95%) |
| --- | --- | --- | --- | --- | --- | --- |

Comment:

**Q6.5: A missense mutation** other than a nonsense mutation [defined as mutation that alters the amino acid encoded by the affected codon] **in *pepQ* gene confer resistance to BDQ.**

| Rarely  (<5%) | Occasionally     (5-24%) | Sometimes  (25-49%) | Frequently  (50-74%) | Frequently  (50-74%) | Very frequently     (75-94%) | Almost always (>=95%) |
| --- | --- | --- | --- | --- | --- | --- |

Comment:

**Q6.6: A homoplasic mutation** [defined as a variant that has arisen multiple times independently] **in *pepQ* is a likely signal for positive selection and confer resistance to BDQ.**

| Rarely  (<5%) | Occasionally     (5-24%) | Sometimes  (25-49%) | Frequently  (50-74%) | Frequently  (50-74%) | Very frequently     (75-94%) | Almost always (>=95%) |
| --- | --- | --- | --- | --- | --- | --- |

Comment:

**Q7** Do you believe that a mutation in ***Rv1979c*** can confer resistance to BDQ?

Yes	No	*If* ***yes****, display sub-questions*

**Q 7.1:** A **synonymous mutation** [defined as mutation that does not alter the amino acid encoded by the affected codon] **in *Rv1979c*** **is neutral** [defined by susceptible phenotype to BDQ]

| Rarely  (<5%) | Occasionally     (5-24%) | Sometimes  (25-49%) | Frequently  (50-74%) | Frequently  (50-74%) | Very frequently     (75-94%) | Almost always (>=95%) |
| --- | --- | --- | --- | --- | --- | --- |

Comment:

**Q 7.2: A** **nonsense mutation** [also called a premature stop codon and defined as a substitution of a single base pair that alters the DNA sequence and leads to the production of a shortened protein] **in *Rv1979c* is a loss of function mutation and confer resistance to BDQ.**

| Rarely  (<5%) | Occasionally     (5-24%) | Sometimes  (25-49%) | Frequently  (50-74%) | Frequently  (50-74%) | Very frequently     (75-94%) | Almost always (>=95%) |
| --- | --- | --- | --- | --- | --- | --- |

Comment:

**Q 7.3: A frameshift indel** [defined as an insertion or deletion of a number-not divisible by three- of nucleotides in a DNA sequence that affects the reading frame of the gene and results in a different translation compared to the wild type] **in *Rv1979c* is a loss of function mutation and confer resistance to BDQ.**

| Rarely  (<5%) | Occasionally     (5-24%) | Sometimes  (25-49%) | Frequently  (50-74%) | Frequently  (50-74%) | Very frequently     (75-94%) | Almost always (>=95%) |
| --- | --- | --- | --- | --- | --- | --- |

Comment:

**Q 7.4: An** **in-frame indel** [defined as an insertion or deletion of a number - a multiple of three - of nucleotides in a DNA sequence which does not shift the reading frame downstream] **in** ***Rv1979c* confer resistance to BDQ.**

| Rarely  (<5%) | Occasionally     (5-24%) | Sometimes  (25-49%) | Frequently  (50-74%) | Frequently  (50-74%) | Very frequently     (75-94%) | Almost always (>=95%) |
| --- | --- | --- | --- | --- | --- | --- |

Comment:

**Q 7.5:** A **missense mutation** other than a nonsense mutation [defined as mutation that alters the amino acid encoded by the affected codon] **in *Rv1979c* confer resistance to BDQ.**

| Rarely  (<5%) | Occasionally     (5-24%) | Sometimes  (25-49%) | Frequently  (50-74%) | Frequently  (50-74%) | Very frequently     (75-94%) | Almost always (>=95%) |
| --- | --- | --- | --- | --- | --- | --- |

Comment:

**Q 7.6: A homoplasic mutation** [defined as a variant that has arisen multiple times independently] **in *Rv1979c* is a likely signal for positive selection and confer resistance to BDQ.**

| Rarely  (<5%) | Occasionally     (5-24%) | Sometimes  (25-49%) | Frequently  (50-74%) | Frequently  (50-74%) | Very frequently     (75-94%) | Almost always (>=95%) |
| --- | --- | --- | --- | --- | --- | --- |

Comment:

**Q 8: On average, how sure were you about the answer you gave to Q 1-9**

- very sure
- relatively sure
- Not so sure
- Not sure at all

***Three questions about you (Q 9-11)***

**Q 9:** what describes you best?

- Academic researcher – mainly laboratory research
- Academic researcher – mainly clinical research
- Academic researcher – mainly epidemiological and public health research
- Clinician

**Q 10:** how many years have you been involved in TB research or TB care?

- < 1 year
- 1 to 5 years
- 6 to 10 years
- 11 to 15 years
- > 15 years

**Q 11:** where do you live?

- Low-income country
- Middle-income country
- High-income country
